# Nested circuits mediate the decision to vocalize

**DOI:** 10.1101/2022.12.14.520381

**Authors:** Shuyun Xiao, Valerie Michael, Richard Mooney

## Abstract

Vocalizations facilitate mating and social affiliation, but may also inadvertently alert predators and rivals. Consequently, the decision to vocalize depends on brain circuits that can weigh and compare these potential benefits and risks. Male mice produce ultrasonic vocalizations (USVs) during courtship to facilitate mating and female mice produce USVs to promote social affiliation with each other. Previously we showed that a specialized set of neurons in the midbrain periaqueductal gray (PAG) are an obligatory gate for USV production in both male and female mice, and that both PAG-USV neurons and USVs can be switched on by their inputs from the preoptic area (POA) of the hypothalamus and switched off by their inputs from neurons on the border between the central and medial amygdala (Amg_C/M-PAG_ neurons) (Michael et al., 2020). Here we show that the USV-suppressing Amg_C/M-PAG_ neurons are strongly activated by predator cues or during social contexts that suppress USV production in male and female mice. Furthermore, a subset of USV-promoting POA neurons that provide input to the PAG-USV region also extend axon collaterals to the amygdala, where they directly inhibit Amg_C/M-PAG_ neurons. Accordingly, Amg_C/M-PAG_ neurons, along with POA_PAG_ and PAG-USV neurons, form a nested hierarchical circuit in which environmental and social information converges to influence the decision to vocalize.

## Introduction

Vocalizations help foster mating and social bonding in most mammals, including rodents (Seyfarth & Cheney, 2010). Therefore, the brain circuits that promote vocalization must respond to salient sexual and social cues generated by receptive conspecifics. However, while vocalizations are important for mating and social affiliation, vocalizing in the wrong context can increase the risk of attracting the attention of eavesdropping predators or social rivals (Akre et al., 2011; Roberts et al., 2007). Therefore, the neural circuits that regulate vocalization must also be able to respond to these threats and weigh them against the potential benefits of mating and social affiliation. How these various and often opposing types of sexual, social and environmental factors are integrated in the brain to influence the decision to vocalize remains unclear. Here we sought to explore the idea that vocal-suppressing neurons in the amygdala influence the decision to vocalize by encoding information about predators and threatening rivals and weighing this against information about potential social partners and mates.

Ultrasonic vocalizations (USVs) are produced by male mice during courtship to facilitate mating and by female mice towards novel female conspecifics, where they are thought to promote social affiliation (Chabout et al., 2015; Maggio & Whitney, 1985; Neunuebel et al., 2015; Portfors, 2007; Warren et al., 2018; Warren et al., 2020; Zhao et al., 2021). As with the production of affiliative vocalizations in other mammals, USV production in mice is enhanced by certain social factors, including the presence of suitable social and sexual partners (Chabout et al., 2015; Zhao et al., 2021). Context-dependent vocal suppression is also a hallmark of the appropriate regulation of USVs during courtship. For example, female mice lacking Trpc2, an ion channel necessary for sensory transduction in the vomeronasal organ, produce USVs at high frequencies in the presence of males, a context in which female mice normally produce few USVs, suggesting that male pheromones normally act to suppress female USV production (Kimchi et al., 2007).

The recent identification of neurons in the midbrain periaqueductal gray (PAG) that serve as an obligatory gate for USV production (PAG-USV neurons) provides the potential for understanding the circuit logic underlying a mouse’s decision to vocalize (Tschida et al., 2019). Notably, a GABAergic projection from the hypothalamic preoptic area (POA) to the PAG drives USV production in male and female mice through disynaptic disinhibition of PAG-USV neurons (Chen et al., 2021; Michael et al., 2020). Conversely, a subpopulation of GABAergic neurons on the border of the central and medial amygdala (Amg_C/M-PAG_ neurons) directly inhibits PAG-USV neurons, and optogenetically activating these neurons in male mice suppresses their courtship USVs (Michael et al., 2020). Such convergent and opponent circuit architecture points to the PAG as one site where vocal-promoting signals could be weighed against vocal-suppressing signals to influence the decision to vocalize. However, while POA neurons that express Estrogen receptor alpha (Esr1), which is a molecular marker of POA neurons that innervate the PAG-USV region (Chen et al., 2021; Michael et al., 2020), are strongly activated in male mice during female courtship (Chen et al., 2021), the extent to which Amg_C/M-PAG_ neurons are excited by predators or social rivals remains unknown. Thus, one question we sought to answer is whether Amg_C/M-PAG_ neurons are activated by predator cues or during social contexts that typically suppress USV production in male and female mice.

Moreover, carefully weighing the potential costs and benefits of vocalization is likely to involve integration at other sites beyond the PAG. Indeed, some POA neurons that project to the PAG-USV region also project to the region of the amygdala that contains Amg_C/M-PAG_ neurons (Michael et al., 2020), raising the possibility that Amg_C/M-PAG_ neurons integrate USV-promoting as well as USV-suppressing information. However, whether POA neurons that provide input to the Amg_C/M_ are active during USV-promoting social encounters and whether activity in this subset of POA neurons is sufficient to drive USV production in the absence of social cues remain untested. Lastly, the physiological properties of synaptic inputs from the POA onto Amg_C/M-PAG_ neurons are unknown, but these properties are important to understand whether the Amg_C/M_ balances competing types of information to determine whether or not a mouse will vocalize.

Here we investigated the role of Amg_C/M-PAG_ neurons and their inputs from the POA in regulating USV production using in vivo fiber photometry of identified cell groups, anatomical and functional circuit mapping, and functional manipulations of neuronal activity in socially interacting and isolated mice. These experiments reveal that Amg_C/M-PAG_ neurons in both male and female mice are strongly activated by cues or contexts that suppress USV production, and that optogenetically activating Amg_C/M-PAG_ neurons selectively suppresses USV production in both male and female mice across a variety of affiliative social contexts in which they typically vocalize. Furthermore, we found that Amg_C/M-PAG_ neurons receive monosynaptic inhibitory input from POA neurons that also project to the PAG, that these inhibitory inputs are active in USV-promoting social contexts, and that optogenetic activation of POA cell bodies that make divergent axonal projections to the amygdala and PAG is sufficient to elicit USV production in socially isolated mice. Together these experiments reveal a nested hierarchical circuit in which environmental and social information is integrated and weighed at multiple processing steps to influence the decision to vocalize.

## Results

### Amg_C/M-PAG_ neurons are active in response to threatening stimuli

Given that optogenetically activating Amg_C/M-PAG_ neurons suppresses USV production in male mice during courtship encounters with females, we reasoned that Amg_C/M-PAG_ neurons would be highly active during threatening contexts in which vocalization is potentially risky for the sender and thus likely to be suppressed. First, we tested how USV production during social encounters was affected by the synthetic fox urine odor 2-methyl-2-thiazoline (2MT), a predator odorant that is highly aversive to mice (Day et al., 2004; Lin et al., 2006; Root et al., 2014). Specifically, we measured the number of USVs produced by male or previously isolated female (see following section) mice during a 5-minute encounter with a novel female conspecific in the presence or absence of 2MT. We found that both male and female mice produced significantly fewer USVs when encounters with a novel female occurred in the presence of 2MT compared to controls (Fig. 1A).

**Figure 1:**
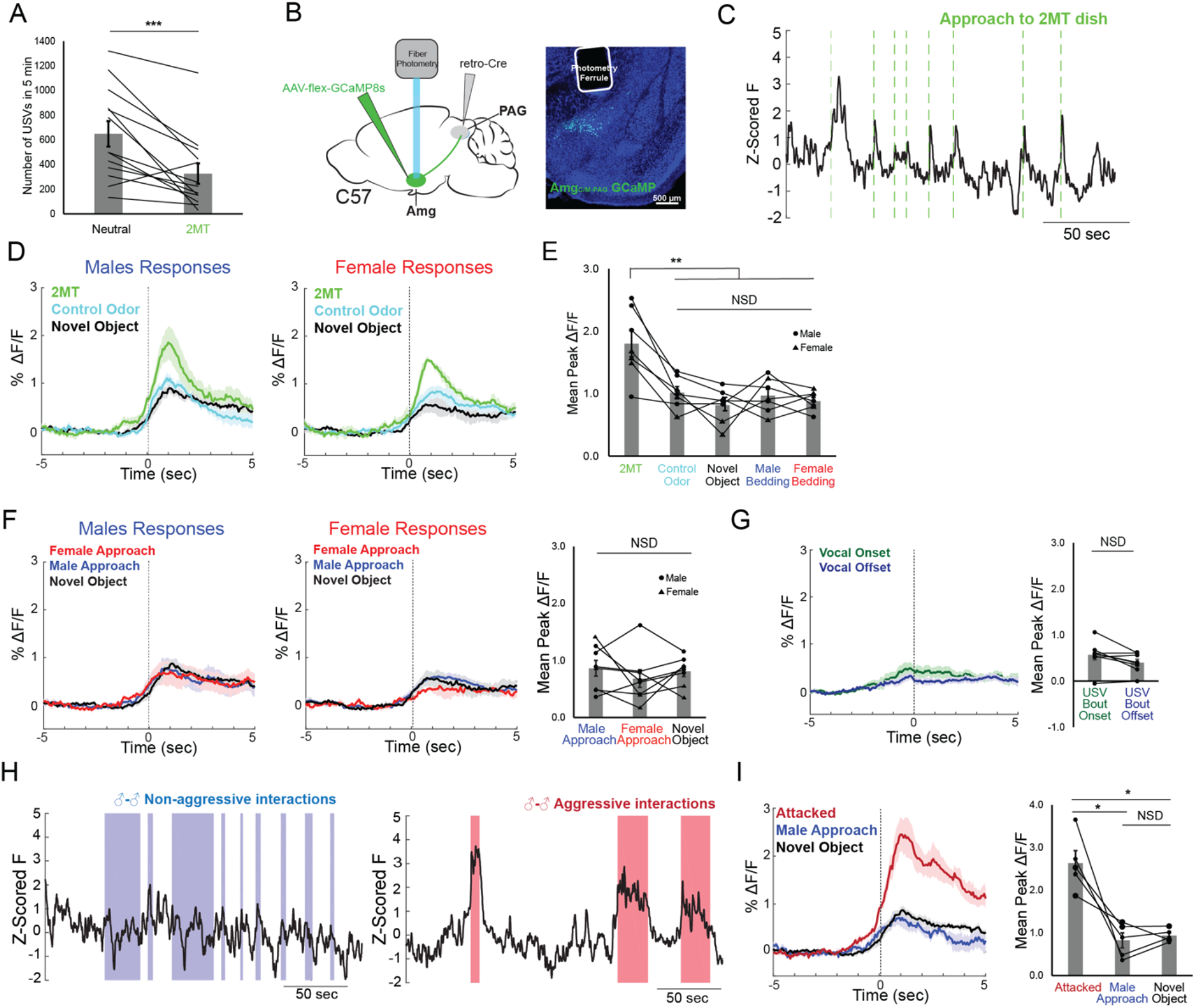
Amg_C/M-PAG_ neurons are active in response to threatening stimuli. (A) The number of USVs produced by male or previously isolated female mice during a 5-minute encounter with a novel female conspecific in the presence or absence of 2MT. (N=13, 11 males, 2 females; p < 0.001; paired *t* test). (B) (Left) Viral strategy for photometry recording of Amg_C/M-PAG_ neurons. (Right) Confocal image showing photometry ferrule tract and GCaMP8s expression in Amg_C/M-PAG_ neurons. (C) Calcium signal in Amg_C/M-PAG_ neurons as a male mouse approached and investigated a dish containing 2MT. Dashed green lines indicate approach onset. (D) Average Amg_C/M-PAG_ activity in males (N=4) and females (N=3) as the animal approached and investigated a dish containing either 2MT, a control odor (ethyl tiglate), or a novel plastic “toy.” (E) Mean peak Amg_C/M-PAG_ activity when approaching a dish containing either 2MT, a control odor (ethyl tiglate) or a novel plastic “toy” or the soiled bedding of male or female conspecifics (N=7, 4 males, 3 females; one-way ANOVA followed by post-hoc pairwise Tukey’s HSD tests). Error bars represent S.E.M. (F) Average (left) and mean peak (right) Amg_C/M-PAG_ activity in males (N=5) and females (N=3) during non-aggressive social encounters with male or female conspecifics, or with a mouse’s investigation of a novel object (one-way ANOVA followed by post-hoc pairwise Tukey’s HSD tests). Error bars represent S.E.M. (G) Average (left) and mean peak (right) Amg_C/M-PAG_ activity aligned to vocal onset and offset (N=5 males; paired *t* test). (H) Calcium signal in Amg_C/M-PAG_ neurons during non-aggressive (left) and aggressive (right) male-male interactions. (I) Average (left) and mean peak (right) Amg_C/M-PAG_ activity in males (N=5) during attack, non-aggressive social encounters with male conspecifics, or with a mouse’s investigation of a novel object (one-way ANOVA followed by post-hoc pairwise Tukey’s HSD tests). Error bars represent S.E.M.

Next, we used an intersectional viral strategy to express the calcium indicator GCaMP8s in Amg_C/M-PAG_ neurons (Fig. 1B) and used fiber photometry to measure calcium signals in these neurons in male and female mice as they investigated a dish containing either 2MT, a control odor (ethyl tiglate), a novel plastic “toy,” or the soiled bedding of male or female conspecifics.

In both sexes, calcium signals in Amg_C/M-PAG_ neurons were elevated above baseline when the animal approached and investigated dishes containing any of these items, but they were the highest when investigating the dish containing 2MT (Fig. 1C-E). The overall pattern of responsiveness to these various stimuli was similar between male and female mice (Fig. 1E; Fig. S1A). Furthermore, the calcium increase evoked by 2MT was not linked to subsequent bouts of freezing, suggesting that activity in these neurons was driven by detection of the odorant rather than the subsequent behavioral response (Fig. S1B). Therefore, an innately aversive predator odorant suppresses USV production and strongly activates USV-suppressing Amg_C/M-PAG_ neurons in both male and female mice.

We then tested how Amg_C/M-PAG_ neuron activity was affected during social interactions with other conspecifics that can either promote or suppress USV production. We used fiber photometry to measure calcium signals of Amg_C/M-PAG_ neurons in freely behaving male and female mice during 5-minute encounters with novel male and female conspecifics. These encounters resulted in non-aggressive interactions when USV production was more likely to occur or aggressive interactions (i.e., fights between two males) when USV production is absent (we confirmed that males did not produce USVs during fights) (Morgret & Dengerink, 1972; Lahvis et al., 2012). In both sexes, we observed similar small but statistically insignificant increases in Amg_C/M-PAG_ neuronal activity when calcium signals were aligned to the onsets of non-aggressive social encounters with male or female conspecifics, or with a mouse’s investigation of a novel object (Fig. 1F; Fig. S1A). Furthermore, calcium signals of Amg_C/M-PAG_ neurons did not change significantly at the onset or offset of USV bouts, suggesting that Amg_C/M-PAG_ neurons are not simply turning off USV bouts during non-aggressive social encounters (Fig. 1G). In contrast, Amg_C/M-PAG_ neuronal calcium signals increased markedly during aggressive interactions between two males that included attacks (Fig. 1H-I). Although many of these epochs included aggressive behavior from both males, Amg_C/M-PAG_ neuronal calcium signals were larger in the male being attacked (Fig. S1C). In summary, Amg_C/M-PAG_ neurons exhibit little or no increase in activity during affiliative social encounters in which USVs frequently occur but are highly activated during aggressive social encounters when USVs are not typically produced. These observations support the idea that Amg_C/M-PAG_ neurons actively suppress USV production in the presence of threatening stimuli.

### Activating PAG-projecting Amg_C/M_ neurons suppresses female USV production

While we previously showed that optogenetic activation of Amg_C/M-PAG_ neurons transiently suppressed courtship USVs in male mice without disrupting non-vocal social behavior (Michael et al., 2020), it remained an open question as to whether these neurons also suppress USV production in female mice. To resolve this issue, we adopted a recently developed behavioral protocol that can reliably elicit robust USV production from female mice (Zhao et al., 2021). In order to elicit high numbers of USVs, female mice were socially isolated for at least three weeks and then, on the day of testing, a novel female conspecific was introduced to the socially isolated experimental animal’s home cage (Fig. 2A). We observed that while group-housed female mice produce few USVs in response to a novel female conspecific, previously isolated female mice produced significantly more USVs per minute, more USV bouts per minute, and longer USV bouts, and in fact vocalized at rates comparable to isolated male mice during female presentation (Fig. 2A). This finding confirms the recent study by Zhao et al. (2021), and provided us with a useful entry point for addressing whether Amg_C/M-PAG_ neurons suppress USV production in female mice.

**Figure 2:**
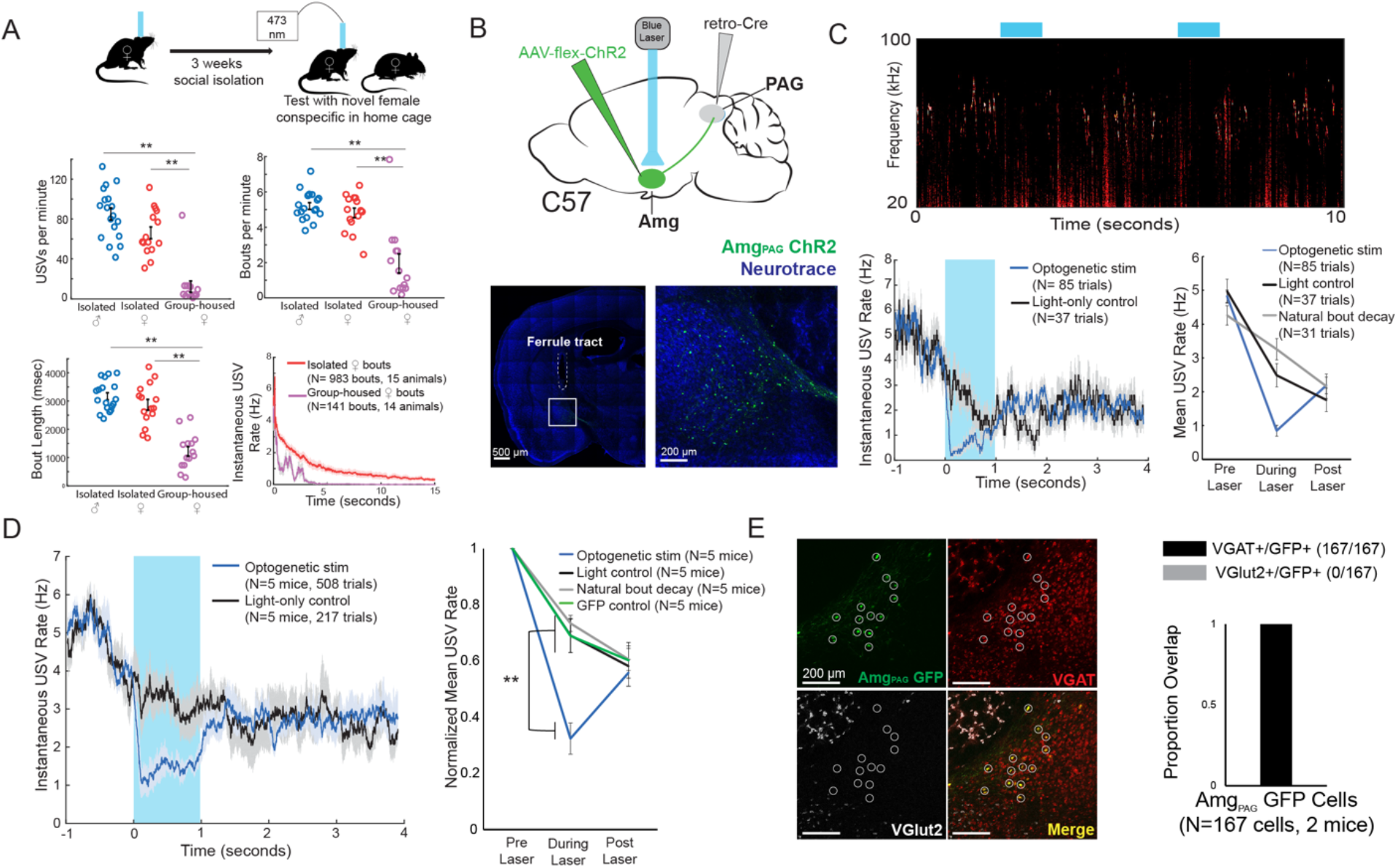
Activation of Amg_C/M-PAG_ neurons suppresses vocalizations in female mice. (A) (Top) Behavioral paradigm for inducing USV production from female mice. (Bottom) Isolated female mice had similar numbers of USVs produced per minute, number of USV bouts produced per minute, and bout lengths when compared to isolated male mice, while group-housed females produced significantly fewer USVs per minute and significantly fewer and shorter bouts (N=18 isolated males, 15 isolated females, and 14 group-housed females, one-way independent sample ANOVAs found a significant effect of condition on USVs/min, USV bouts/min, and USVs/bout, post hoc pairwise Tukey’s HSD tests found that for all three measures *p*<0.01 for isolated males versus group housed females, *p*<0.01 for isolated females versus group housed males, and NSD between isolated males and isolated females). Error bars represent S.E.M. (B) (Top) Viral strategy for optogenetic activation of Amg_C/M-PAG_ neurons in female mice (performed in N=5 females). (Bottom) Confocal image of representative Amg_C/M-PAG_ cell body labeling and ferrule placement achieved with this strategy. (C) (Top) Example spectrogram showing a representative set of trials in which activation of Amg_C/M-PAG_ neurons suppresses ongoing female USV production. (Bottom Left) Example traces of the USV rate during optogenetic stimulation vs a light-control condition for all trials (N=85 optogenetic trials and 37 light-only control trials) for a single representative animal. Error shading above and below the mean represents S.E.M. (Bottom Right) Quantification of the mean USV rate 1 second before laser stimulation, during a 1 second laser stimulation, and 1 second after laser stimulation (blue) compared to light-only control trials (black) and to the natural decay of vocal bouts with no stimulation (gray) for one representative animal. Error bars represent S.E.M. (D) (Left) Average USV rate traces during optogenetic stimulation and a light-only control for all animals (N=5 females). Error shading above and below the mean represents S.E.M. (Right) Group averages of the mean USV rate 1 second before laser stimulation, during a 1 second laser stimulation, and 1 second after laser stimulation (blue) compared to light-only control trials (black) and to the natural decay of vocal bouts with no stimulation (gray) for all optogenetic stimulation animals (N=5 females) and for a separate group of GFP control animals (N=5 females, green). Data for each mouse were normalized by dividing the mean USV rate pre, during, and post laser stimulation by the mean USV rate for the pre-laser period (*p*<0.01 for differences between ChR2 group vs. control groups during laser time; two-way ANOVA with repeated measures on one factor, *p*<0.001 for interaction between group and time, followed by post-hoc pairwise Tukey’s HSD tests; *p*>0.05 for differences between groups in post-laser period). Error bars represent S.E.M. (E) (Left) Representative confocal image of *in situ* hybridization performed on Amg_C/M-PAG_ neurons (labeled with GFP, shown in green), showing overlap with expression of VGAT (red) and VGlut2 (white). (Right) Quantification of overlap of GFP-labeled Amg_C/M-PAG_ neurons with VGAT and VGlut2 (N=167 from 2 female mice).

To this end, we used an intersectional viral strategy to selectively express channelrhodopsin (ChR2) in Amg_C/M-PAG_ neurons of female mice (Fig. 2B). We found that this strategy labels neurons in the Amg_C/M_ but not the CeA of female mice, similar to the pattern previously observed with this strategy in male mice (Fig. 2B; see also Fig. 4A in Michael et al, 2020). Using the social isolation protocol, we were able to elicit high numbers of USVs routinely in 5 of 7 Amg_C/M-PAG_-ChR2 female mice as they investigated a novel female partner. Optogenetic stimulation of Amg_C/M-PAG_ neurons in freely vocalizing female mice immediately suppressed USV production, and this suppression persisted throughout the duration of the laser pulse (Fig. 2C-D, N=5 of 5 female mice). The optogenetically-evoked decrease in USV rate was significant when compared to a light-only control condition in which the same mice were connected to a dummy ferrule that shone blue light above their heads, a control condition in which the same mice did not receive laser stimulation and USV rates decreased naturally, and to a separate group of female mice in which GFP was expressed in Amg_C/M-PAG_ neurons and that were subjected to optical fiber illumination in the Amg (Fig. 2D, N=5 female mice in the optogenetic condition, light control condition, and natural bout decay condition and N=5 female mice in the GFP control condition, *p*<0.01 for differences between ChR2 group vs. control groups during laser time; two-way ANOVA with repeated measures on one factor, *p*<0.01 for interaction between group and time, followed by post-hoc pairwise Tukey’s HSD tests). After laser offset, the USV rate in the Amg_C/M-PAG_-ChR2 female mice rebounded to a level comparable to that in all control conditions (*p*>0.05 for differences between groups in post-laser period; Fig. 2C-D). In male mice, USV-suppressing Amg_C/M-PAG_ neurons are predominantly GABAergic (Michael et al., 2020). We used in situ hybridization to establish that 100% of Amg_C/M-PAG_ cells in female mice express VGAT, suggesting that they mediate USV suppression by directly inhibiting PAG-USV neurons (N=167/167 neurons in 2 female mice were VGAT+ and 0/167 were VGlut2+; Fig. 2E), as described in males (Michael et al., 2020). In summary, Amg_C/M-PAG_ neurons are GABAergic and function to suppress vocalizations in female mice during social encounters with other females, similar to the manner in which these neurons suppress courtship USVs in male mice. Therefore, although female and male mice produce USVs in different social and sexual contexts, Amg_C/M-PAG_ neurons in both sexes function similarly to suppress USV production.

### Amg_C/M-PAG_ neurons that suppress vocalization express estrogen receptor alpha (Esr1)

Neurons that express estrogen receptor alpha (Esr1) have been implicated in a wide range of sexual and social behaviors including aggression, social recognition, courtship and, notably, vocalization (Chen et al., 2021; Hashikawa et al., 2017; Kudwa et al., 2006; Lee et al., 2014; Michael et al., 2020; Moran et al., 2020). Indeed, several recent studies showed that optogenetically activating Esr1^+^ neurons in the POA was sufficient to elicit long-lasting bouts of USVs in both male and female mice (Chen et al., 2021; Karigo et al., 2021; Michael et al., 2020). Given these observations, we wondered whether Esr1 might also be expressed in Amg_C/M-PAG_ neurons. Consistent with this idea, *in situ* hybridization data from the Allen Brain Atlas shows expression of Esr1 mRNA in the same region of the amygdala as Amg_C/M-PAG_ neurons are found in male mice (Fig. 3A). We therefore tested the hypothesis that Amg_C/M-PAG_ neurons express Esr1 using immunofluorescent Esr1 protein antibody staining. We found that nearly half of GFP-labeled Amg_C/M-PAG_ neurons express Esr1 (Fig. 3B, ∼49%, N=913 cells from 7 mice, 3 males and 4 females; Amg_C/M-PAG_ neurons labeled by injecting PAG with AAV-retro-Cre and amygdala with AAV-flex-GFP, similar to the strategy shown in Fig. 2B). By injecting the amygdala of Esr1-cre mice with flex-tdTomato we further found that Esr1^+^ amygdala cells send axonal projections to the region of caudolateral PAG, the region that contains PAG-USV neurons, and consistent with our observation that a large subset of Amg_C/M-PAG_ neurons are Esr1^+^ (Fig. 3C).

**Figure 3:**
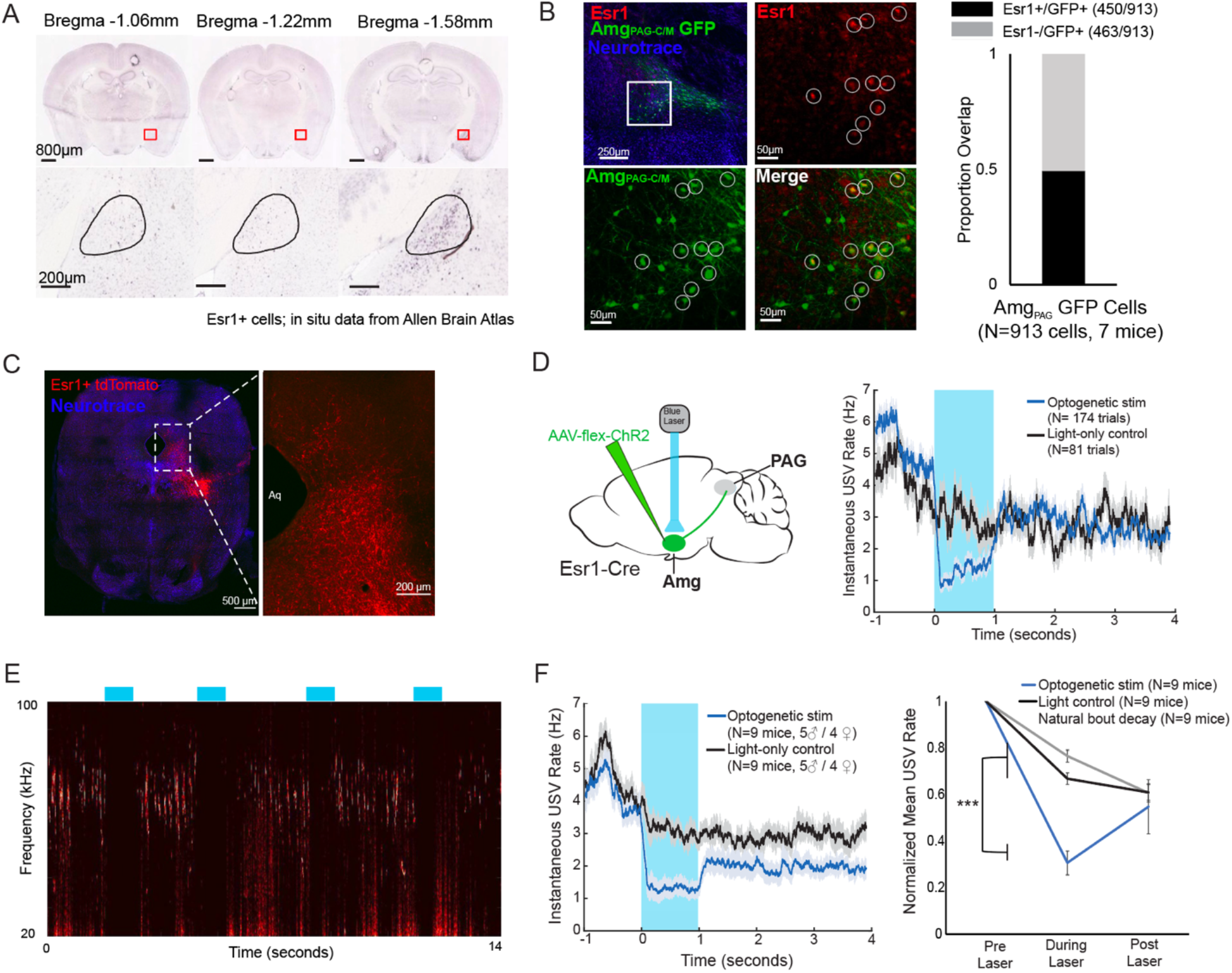
Activation of Esr1+ Amg_C/M_ neurons suppresses vocalizations. (A) *In situ* hybridization data for Esr1 mRNA in wild-type male mouse. (Allen Institute for Brain Science. Allen Mouse Brain Atlas, available from: http://mouse.brain-map.org/experiment/show/79591677). (B) (Left) Representative confocal image of immunofluorescent Esr1 protein antibody staining performed on Amg_C/M-PAG_ neurons (labeled with GFP, shown in green), showing overlap with expression of Esr1 (red). Blue is NeuroTrace. (Right) Quantification showing overlap of GFP-labeled Amg_C/M-PAG_ neurons with Esr1 (N=913 cells from 7 mice; 3 males and 4 females). (C) Axonal projections of Esr1^+^ amygdala cells (labeled by tdTomato) in caudolateral PAG, the region that contains PAG-USV neurons. (D) (Left) Viral strategy used to express ChR2 in Esr1+ Amg_C/M_ neurons. (Right) Example traces of the USV rate during optogenetic stimulation vs a light-control condition for all trials (N=174 optogenetic trials and 62 light-only control trials) for a single representative Esr1-Cre-ChR2 animal. Error shading above and below the mean represents S.E.M. (E) Example spectrogram showing a representative set of trials in which activation of Amg_C/M-PAG_ neurons suppresses ongoing USV production. (F) (Left) Same as Figure 1D for all Esr1-Cre-ChR2 animals (N=9 mice, 5 males and 4 females). Error shading above and below the mean represents S.E.M. (Right) Same as Figure 4D but for all optogenetic stimulation animals (N=9 mice, 5 males and 4 females). Data for each mouse were normalized by dividing the mean USV rate pre, during, and post laser stimulation by the mean USV rate for the pre-laser period (p < 0.001 for differences between optogenetic condition and both control conditions during the laser stimulation period, two-way ANOVA followed by post-hoc pairwise Tukey’s HSD tests). Error bars represent S.E.M.

To test whether activation of Esr1^+^ Amg_C/M_ cells is sufficient to suppress USV production, we injected the Amg_C/M_ of Esr1-Cre male and female mice with a Cre-dependent AAV driving ChR2 expression (Fig. 3D). All mice were single-housed post-surgery. Optogenetic activation of Esr1^+^ Amg_C/M_ cells immediately and reversibly suppressed USV production (N=9 mice; 5 males and 4 females; Fig. 3D-F). As with stimulation of Amg_C/M-PAG_ cells, this suppression was specific to the time period of optogenetic stimulation and did not occur in light-only control conditions or in control conditions in which USV bouts were allowed to decay naturally (Fig. 3F; *p*<0.001 for differences between optogenetic condition and both control conditions during the laser stimulation period, two-way ANOVA followed by post-hoc pairwise Tukey’s HSD tests). Taken together, this evidence suggests that Esr1^+^ Amg_C/M-PAG_ neurons mediate vocal suppression in both male and female mice. Though GABAergic Esr1^+^ neurons in the hypothalamus and glutamatergic Esr1+ neurons in the posterior amygdala have been implicated in mediating sexual behaviors and aggression, to our knowledge this is a novel role for GABAergic Esr1^+^ neurons in the Amg_C/M-PAG_.

### Characterizing monosynaptic inputs to Amg_C/M-PAG_ neurons in male mice

The decision to suppress ongoing vocalizations depends on the integration of external cues with interoceptive information. Given that Amg_C/M-PAG_ neurons suppress vocalizations by directly inhibiting vocal gating neurons in the PAG, they are likely to integrate a wide range of inputs that help regulate this decision. We conducted a series of additional anatomical and functional experiments in male mice to better understand the nature of this integrative process. We first conducted monosynaptically-restricted transsynaptic rabies tracing from Amg_C/M-PAG_ neurons to map their inputs (Fig. 4A). To specifically label inputs to Amg_C/M_ neurons that project to PAG we injected the caudolateral PAG with AAV-retro-Cre and the Amg_C/M_ with Cre-dependent helper viruses that express rabies glycoprotein (G), a protein required for transsynaptic spread (Callaway & Luo, 2015). Two weeks later, we injected the same amygdalar regions with a pseudotyped G-deleted rabies virus, which when complemented by the wild-type rabies G glycoprotein, resulted in expression of GFP in neurons that make synapses onto Amg_C/M-PAG_ neurons (Fig. 4A; N=4 males).

**Figure 4:**
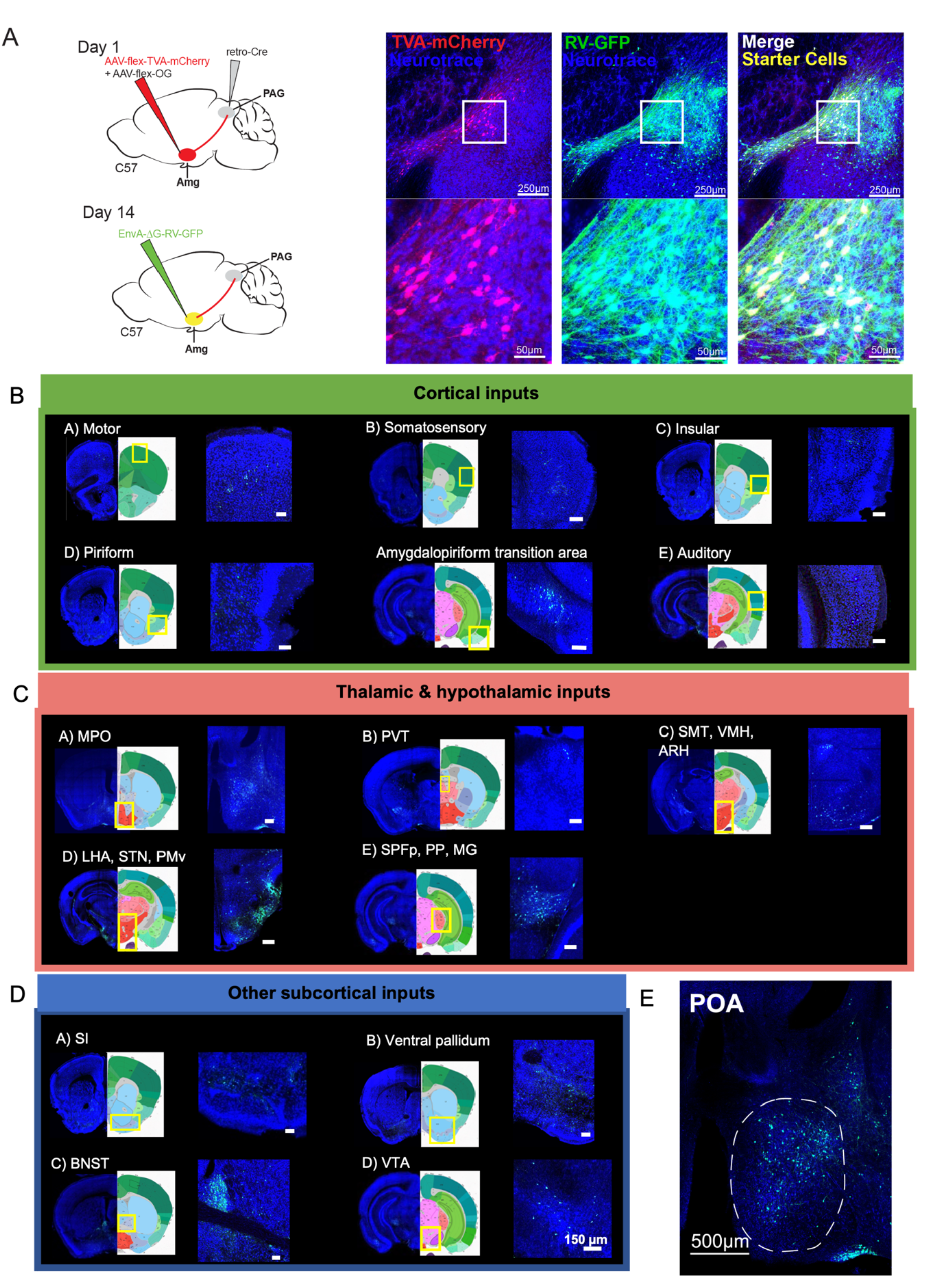
Monosynaptic inputs to Amg_C/M-PAG_ neurons in male mice. (A) (Left) Viral strategy shown for transsynaptic labeling of direct inputs to Amg_C/M-PAG_ neurons (performed in N=4 males). (Right) Confocal images are shown of starter Amg_C/M-PAG_ neurons. (B) – (E) Confocal images are shown of upstream neurons labeled in the cortical and subcortical areas, including the preoptic area of the hypothalamus (POA). Scale bars in B, C, D are 150 μm.

This approach revealed that Amg_C/M-PAG_ neurons in male mice receive monosynaptic input from numerous cortical regions that process a wide variety of sensory and interoceptive information, including the piriform, motor, somatosensory, insular and auditory cortices (Fig. 4B) (Bhattacharjee et al., 2021; Gogolla, 2017; Shipley & Ennis, 1996; Zatorre et al., 2002). We also observed that Amg_C/M-PAG_ neurons receive input from many other forebrain and midbrain regions, including the medial and lateral POA, the bed nucleus of the stria terminalis (BNST), the ventral tegmental area, and the nucleus accumbens shell (Fig. 4C-D), which have been variously implicated in defensive behaviors, sexual-social interactions, reward and satisfaction (Breitfeld et al., 2015; Michael et al., 2020; Paredes, 2003; Salgado & Kaplitt, 2015).

Of particular interest to us was the labeling of cell bodies in the medial POA (Fig. 4E), a region that contains neurons whose activation elicits USVs via a disinhibitory circuit motif within the PAG (Chen et al., 2021; Michael et al., 2020). This anatomical organization raises the possibility that Amg_C/M-PAG_ cells are a cellular locus where USV-suppressing information can be weighed against USV-promoting signals from the POA. Therefore, we focused on characterizing the social and USV-related information that POA axons convey to the Amg_C/M_, and the function of the synapses that POA neurons make on Amg_C/M-PAG_ neurons.

### In male mice, POA inputs to the Amg_C/M_ are active during affiliative social encounters and USV production

To measure the activity of POA inputs to the Amg_C/M_, we used intersectional methods to express GCaMP8s in Esr1+ POA neurons and recorded bulk fluorescence from their axon terminals in the Amg_C/M_ (Fig. 5A; N=5 male Esr1-Cre mice). We then measured the activity of POA axons in the Amg_C/M_ during the male’s social interactions with novel female and male mice, 2MT, or novel plastic toys. In contrast to the activity of Amg_C/M-PAG_ neurons measured in these various conditions (Fig. 1D), POA_Esr1-Amg_ terminal activity showed a large and sustained increase at the onset of social interactions with female mice, a moderate sustained increase at the onset of non-aggressive social interactions with other male mice, and no response at the onset of or during trials where the mouse encountered a dish containing 2MT or a novel plastic toy (Fig. 5B). In addition to the large and sustained increase in activity upon the introduction of a social partner, POA_Esr1-Amg_ terminal activity further increased at the onset of subsequent social interactions and at the onset of individual USV bouts (Fig. 5C-D). However, POA_Esr1-Amg_ terminal activity remained low when aligned to the onsets of the mouse’s investigation of a dish containing 2MT or the onsets of attacks by male conspecifics (Fig. 5E, F). Taken together, these findings support the idea that POA_Esr1-Amg_ axons in the Amg_C/M-PAG_ of male mice are highly active during social encounters in which USVs are produced, but not in situations in which Amg_C/M-PAG_ activity is high and USV production is suppressed. Therefore, in addition to being strongly activated by stimuli or aggressive social interactions that suppress USV production (Fig. 1A, D, I), Amg_C/M-PAG_ neurons receive inputs from the POA that convey information about pro-vocal social contexts and USV production.

**Figure 5:**
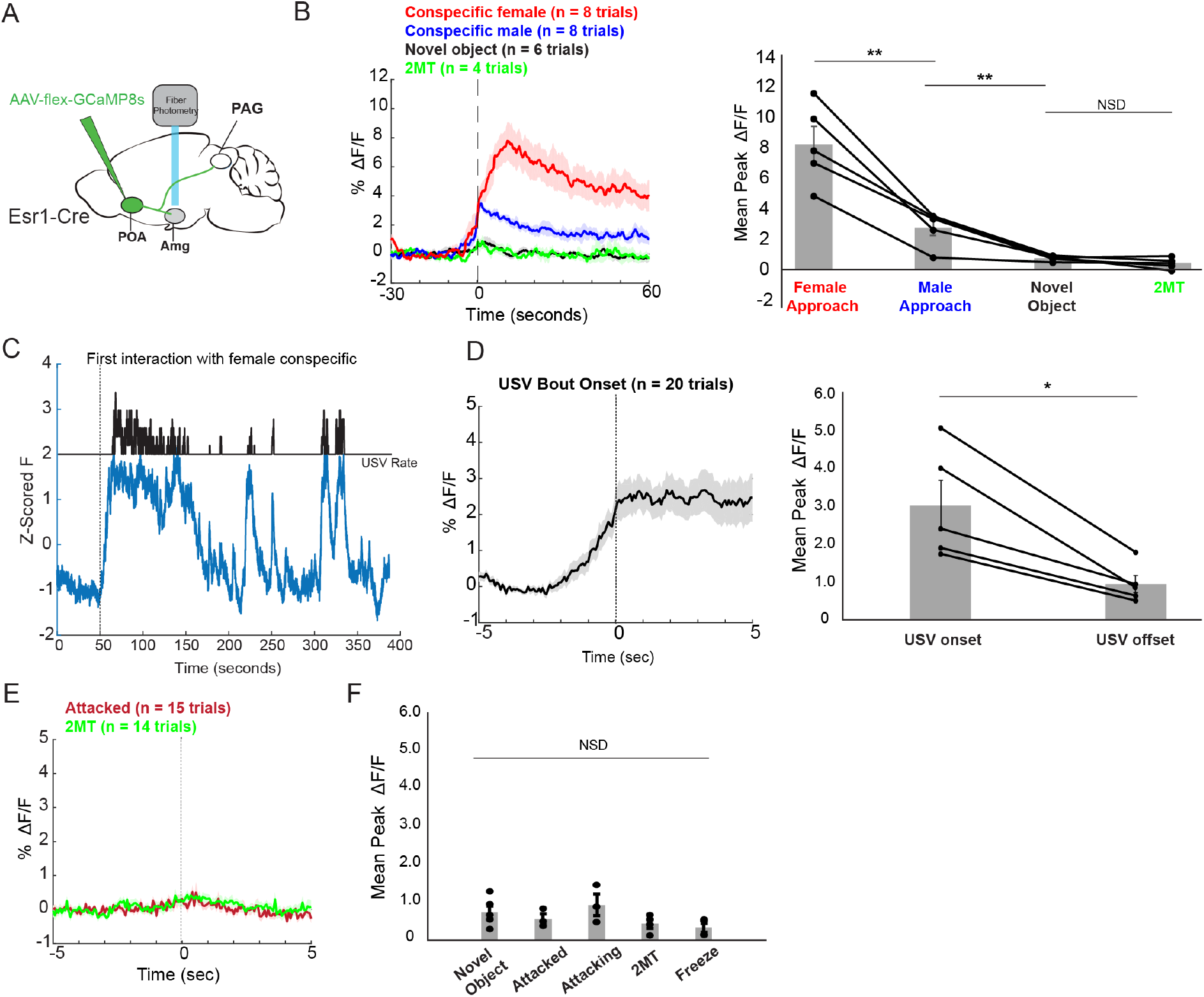
POA inputs to the Amg_C/M_ are active during affiliative social encounters and USV production in male mice. (A) Viral strategy for photometry recording of Esr1+ POA terminal activity in Amg_C/M_ region (POA_Esr1-Amg_ terminal activity). (B) Average POA_Esr1-Amg_ activity in an example male mouse (left) and mean peak POA_Esr1-Amg_ activity from 5 males (right) at the onset of approaching female mice, at the onset of non-aggressive social interactions with other male mice, and at the onset of or during trials where the mouse encountered a dish containing 2MT or a novel plastic toy (N=5 males; one-way ANOVA followed by post-hoc pairwise Tukey’s HSD tests). Error bars represent S.E.M. (C) POA_Esr1-Amg_ terminal activity as a male vocalized to a female. (D) Average POA_Esr1-Amg_ terminal activity in an example male mouse (left) and mean peak POA_Esr1-Amg_ activity from 5 males (right) at USV bout onset (N=5 males; paired *t* test). (E) Average POA_Esr1-Amg_ terminal activity in an example male mouse during aggressive interactions with other males and investigation of a dish containing 2MT. (F) Mean peak POA_Esr1-Amg_ activity from 5 males during aggressive interactions with other males, investigation of novel object or a dish containing 2MT, or during freezing in the presence of 2MT.

### In male mice, POA neurons that provide input to the Amg_C/M_ are Esr1+ and optogenetically activating them is sufficient to promote USV production

These various observations strongly suggest that the POA neurons that provide input to the amygdala are actually USV-promoting neurons. Because activating POA_PAG_ cells that express Esr1 is sufficient to promote USV production (Chen et al., 2021; Karigo et al., 2021; Michael et al., 2020), we first investigated whether POA cells that project to the Amg_C/M_ (POA_AMG_ cells, labeled by intersectional methods; Fig. 6A) are part of this Esr1-expressing subpopulation of POA cells. We found that the majority (∼82%) of GFP-labeled POA_AMG_ cells express Esr1 (Fig. 6B, N=298/362 cells from 2 male mice). We further noted that axon terminals from GFP-labeled POA_AMG_ cells were present in the caudolateral PAG, the same region where PAG-USV neurons are located (Fig. 6C, see also Tschida et al., 2019). To directly test whether POA_AMG_ cells are USV-promoting, we used intersectional viral methods to express ChR2 in POA_AMG_ cells (Fig. 6D). We found that exciting the cell bodies of these ChR2-expressing POA_AMG_ cells with blue light was sufficient to trigger USVs in male mice in the absence of social cues (Fig. 6E-G; N=4 of 6 male mice showed increased USV rates in response to this optogenetic stimulation). Therefore, the POA cells that convey pro-vocal social signals and USV-related signals to the Amg_C/M_ are Esr1+, also project to the caudolateral PAG, and directly promote USV production.

**Figure 6:**
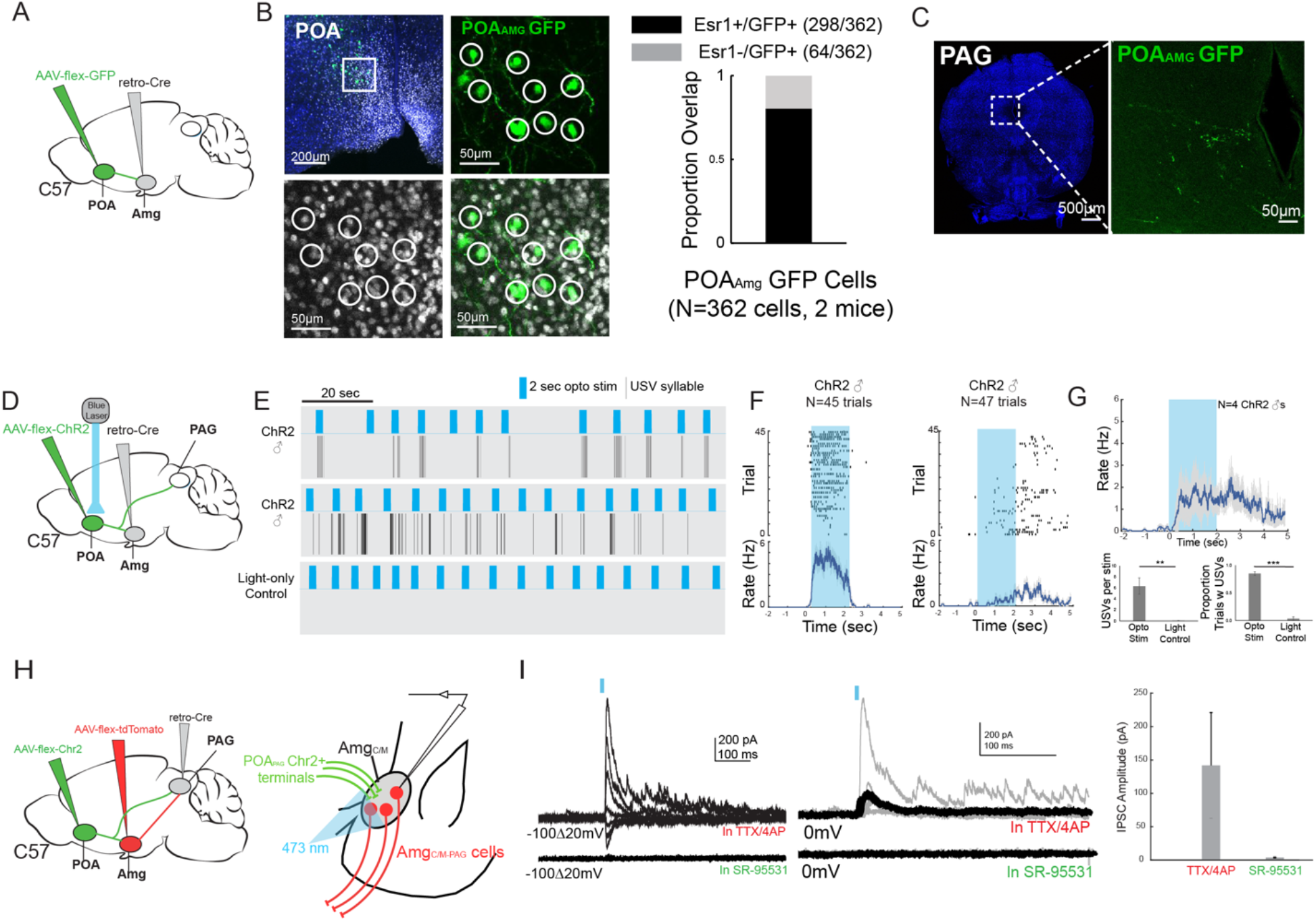
POA_Amg_ cells are Esr1+, project to the PAG, promote USV production, and make inhibitory synapses onto Amg_C/M-PAG_ neurons. (A) Viral strategy for labeling Amg_C/M_-projecting POA neurons. (B) (Left) Confocal images showing GFP-labelled Amg_C/M_-projecting POA neurons and Esr1 staining. (Right) Quantification of Esr1^+^ POA_Amg_ neurons. (C) Axon terminals of GFP-labelled Amg_C/M_-projecting POA neurons in PAG. (D) Viral strategy for optogenetic activation of Amg_C/M_-projecting POA neurons in male mice (N=4). (E) – (G) Optogenetic stimulation of ChR2-expressing POA_AMG_ cells triggered USVs in male mice in the absence of social cues. (H) Viral strategy for in vitro whole-cell voltage clamp recordings from visually identified Amg_C/M-PAG_ neurons while optogenetically activating POA_PAG_ axon terminals in brain slices containing the amygdala. (I) (Left) Light-evoked IPSCs recorded in TTX/4AP (observed in N=9 of 13 td-Tomato-tagged Amg_C/M-PAG_ neurons) were abolished by application of gabazine (N=4 cells also recorded in gabazine). IPSC amplitude refers to the peak of the light-evoked current at 0 mV holding potential. (Right) Mean IPSC Amplitude recorded in TTX/4AP (N=9) and SR-95531 (N=4). Error bars represent S.E.M.

### Vocalization-triggering POA_PAG_ cells make inhibitory synapses onto Amg_C/M-PAG_ neurons

Prior studies show that the vast majority of POA_PAG_ neurons, including those that promote USV production, release the inhibitory neurotransmitter GABA (Chen et al., 2021; Michael et al., 2020). Therefore, our current observations raise the interesting possibility that the Amg_C/M-PAG_ cells that suppress USVs are *directly* inhibited by the POA_PAG_ neurons that promote USV production. To test this hypothesis, we first injected the PAG with AAV-retro-Cre, the POA with AAV-flex-ChR2, and the amygdala with AAV-flex-tdTomato (Fig. 6H). Several weeks later, we performed *in vitro* whole-cell voltage clamp recordings from visually identified Amg_C/M-PAG_ neurons while optogenetically activating POA_PAG_ axon terminals in brain slices containing the amygdala. We visually targeted our recordings to tdTomato^+^ Amg_C/M-PAG_ neurons in the presence of TTX and 4AP to isolate monosynaptic currents. Optogenetically activating POA_PAG_ axon terminals in the amygdala evoked inhibitory postsynaptic currents (IPSCs) in a majority (9/13) of Amg_C/M-PAG_ (td-Tomato+) neurons (Fig. 6I; mean current=141.9 pA at 0mV in TTX/4AP; currents with reversal potentials at -70 mV were identified as IPSCs, and no EPSCs were observed at -70 mV in the presence of TTX and 4AP). Bath application of the GABA_A_ receptor antagonist gabazine SR-95531 abolished these optogenetically-evoked IPSCs in all four cases in which it was applied (Fig. 6I, 4/9 recordings). Therefore, POA_PAG_ cells provide monosynaptic inhibitory input onto Amg_C/M-PAG_ neurons. These various findings support the idea that USV-suppressing Amg_C/M-PAG_ neurons, which are strongly activated by predator cues and social threats, are also directly inhibited by USV-promoting POA neurons.

## Discussion

Here we show that predator cues and aversive social encounters suppress affiliative and courtship USVs produced by female and male mice, and that these cues and contexts directly excite USV-suppressing neurons located at the border of the central and medial amygdala (Amg_C/M_ neurons). Using behavioral methods, we established that 2MT, a component of fox urine suppresses USVs produced by male and female mice, while directly exciting Amg_C/M_ neurons suppresses USV production through their projections to the caudolateral PAG, a region that contains neurons that gate USV production (i.e., PAG-USV neurons) (Tschida et al., 2019). Furthermore, using fiber photometric calcium imaging, we found that these PAG-projecting Amg_C/M_ neurons (Amg_C/M-PAG_ neurons) are strongly excited in the presence of 2MT but not active when the animal subsequently exhibited 2MT-evoked freezing. We also observed that Amg_C/M-PAG_ neurons are excited during aggressive social encounters between male mice, particularly when a male is under attack, a context when we never detected USV production. Thus, this study provides insight into how certain aversive stimuli act centrally in the mouse’s brain to suppress courtship and affiliative vocalizations. Furthermore, we used monosynaptic rabies virus tracing to map an extensive array of cortical and subcortical neurons that provide input onto these Amg_C/M-PAG_ neurons neurons in male mice. These include GABAergic neurons in the medial preoptic area that we and others previously demonstrated promote USV production by disinhibiting PAG-USV neurons (Chen et al., 2021; Michael et al., 2020). Notably, we showed that a subset of GABAergic POA neurons that project to the PAG (POA_PAG_ neurons) also make inhibitory synapses on Amg_C/M-PAG_ neurons, that these POA_PAG_ inputs to Amg_C/M-PAG_ neurons are activated in USV-promoting and USV-producing contexts, and that optogenetically activating POA neurons that provide inputs to the Amg_C/M_ region is sufficient to promote USV production in socially isolated male mice. Consequently, our study advances both Amg_C/M-PAG_ and PAG-USV neurons as loci that weigh competing types of sexual, social and environmental information to regulate the mouse’s decision to vocalize.

While the PAG has long been recognized as an obligatory for vocalizations in a wide range of mammalian species (Jürgens, 2002; Jürgens, 2009), the molecular genetic identification of USV-gating neurons in the mouse has opened the door to a more systematic analysis of the circuitry that underlies the mouse’s decision to vocalize (Michael et al., 2020; Tschida et al., 2019). Indeed, our recent prior study exploited this advance to establish that PAG-USV neurons receive USV-promoting inputs from the POA and USV-suppressing inputs from Amg_C/M-PAG_ neurons (Michael et al., 2020). Consequently, establishing that the Amg-PAG pathway responds to predator odorants and, in the male, threatening social encounters, advances the idea that the PAG is one site where vocal promoting and vocal suppressing information can be weighed to influence the decision to vocalize. As PAG-USV neurons receive highly convergent input from neurons located in a wide variety of brain regions, the PAG presumably integrates additional environmental, social, sexual and interoceptive information in the process of gating vocal production.

Instead of the PAG serving as the sole site where such competing information is weighed, the present study emphasizes that the circuitry that underlies the decision to vocalize is distributed across two brain regions, including the vocal-suppressing neurons in the amygdala as well as USV-gating neurons in the PAG. In particular, here we showed that USV-suppressing Amg_C/M-PAG_ neurons are directly inhibited by USV-promoting neurons in the POA, while a prior study showed that Amg_C/M-PAG_ neurons do not project back into the USV-promoting region of the POA (Michael et al., 2020). Although Amg_C/M-PAG_ neurons may alter POA activity through indirect pathways, the asymmetry of this circuit supports a logic where the reproductive and social benefits of vocalizing may outweigh the potential costs of advertising one’s location to predators and aggressive social rivals. Along these lines, we found 2MT did not totally suppress USV output, implying that USV-promoting social factors can override the USV-suppressing effects of this aversive predator cue.

While male and female mice produce USVs in different amounts and in different social and reproductive contexts, this and prior studies emphasize that many components of the circuitry that regulates USV production are qualitatively similar in the two sexes (Chen et al., 2021; Karigo et al., 2021; Michael et al., 2020; Tschida et al., 2019). This may in turn implicate sexually dimorphic features at earlier stages of processing, particularly of olfactory cues, as a major determinant of sexually dimorphic vocal behavior in mice. Lastly, the present study shows that USV-suppressing Amg_C/M-PAG_ neurons express Esr1, which also distinguishes USV-promoting neurons in the POA (Chen et al., 2021; Michael et al., 2020). Therefore, Esr1 may serve as a more widespread marker of the neural circuits that regulate USV production in both male and female mice. In summary, the present study provides insight into how environmental and social factors are encoded and weighed at several levels of a hierarchically organized circuit to regulate affiliative vocalizations in male and female mice. Given the conserved nature of many of the neural components examined here, a similar nested circuit design may play an important role in regulating vocalizations in other mammals, including humans.

## Materials and Methods

### CONTACT FOR REAGENT AND RESOURCE SHARING

Further information and requests for resources and reagents should be directed to the corresponding authors, Richard Mooney.

### EXPERIMENTAL MODELS AND SUBJECT DETAILS

#### Animal statement

All experiments were conducted according to protocols approved by the Duke University Institutional Animal Care and Use Committee.

#### Animals

For optogenetic activation, fiber photometry, and transsynaptic tracing experiments, the following mouse lines from Jackson labs were used: C57 (000664) and Esr1-Cre (017911). For whole-cell recording experiments C57 mice were used (000664).

### METHOD DETAILS

#### Viruses

The following viruses and were used: AAV2/1-hSyn-flex-ChR2-eYFP (Addgene), AAV-pgk-retro-Cre (Addgene), AAV-flex-GFP (Addgene), AAV-flex-tdTomato (Addgene), AAV-flex-GCamp8s (Addgene), and AAV-flex-oG (Duke Viral Vector Core). EnvA-ΔG-RV-GFP and AAV-flex-TVA-mCherry were produced in house as previously described (Rodriguez et al., 2017; Sakurai et al., 2016; Tschida et al., 2019, Michael et al., 2020). The final injection coordinates were as follows: POA, AP=0.14 mm, ML= 0.3 mm, DV= 5.5 mm; Amg_C/M_, AP=-1.5 mm, ML= 2.3 mm, DV= 4.6 mm; PAG, AP=-4.7 mm, ML= 0.7 mm, DV= 1.75 mm. Viruses were pressure-injected with a Nanoject II (Drummond) at a rate of 4.6 nL every 15s.

#### In vivo optogenetic stimulation

For optogenetic stimulation of Amg_C/M-PAG_ cells in female mice, the caudolateral PAG of C57 female mice was injected with AAV-pgk-retro-Cre (100 nL) and the Amg_C/M_ was injected with AAV-flex-ChR2 (100 nL) and the optogenetic ferrule was placed above the Amg_C/M_ cell bodies. For optogenetic stimulation of Amg_C/M_ Esr1+ neurons the Amg_C/M_ of Esr1-cre male and female mouse was injected with AAV-flex-ChR2 (100 nL) and the optogenetic ferrule was placed above the Amg_C/M_ cell bodies. For optogenetic stimulation of POA_Amg_ cells the Amg_C/M_ of C57 male and female mice was injected with AAV-pgk-retro-Cre (100 nL) and the medial POA was injected with AAV-flex-ChR2 (100 nL) and the optogenetic ferrule was placed above the POA_Amg_ cells bodies.

Commercially available (RWD) optogenetic ferrules were implanted in the same surgeries as viral injection just above target brain locations and were fixed to the skull using Metabond (Parkell). Neurons were optogenetically activated with illumination from a 473 nm laser (3-15 mW) at 10-20 Hz (50 ms pulses, 2-10s total) or with phasic laser pulses (1-2s duration). Laser stimuli were driven by computer-controlled voltage pulses (Spike 7, CED). For stimulation of Amg_C/M-PAG_ neurons, Amg_C/M_ Esr1+ neurons, the laser was manually triggered each time the mouse began a bout of vocalization towards a social partner that lasted several seconds. For stimulation of POA_Amg_ neurons stimulation was triggered manually at regular intervals while the animal was alone.

#### USV recording and analysis

as described in Michael et al. (2021).

#### Transsynaptic tracing from Amg_C/M-PAG_ neurons

For selective transsynaptic tracing from Amg_C/M-PAG_ neurons with viruses, the caudolateral PAG was injected with AAV-pgk-retro-Cre and the Amg_C/M_ was injected with a 1:1 mixture of AAV-flex-TVA-mCherry and AAV-flex-oG (total volume of 100 nL). After a wait time of 10-14 days, the Amg_C/M_ was then injected with EnvA-ΔG-RV-GFP (100 nL, diluted 1:5), and animals were sacrificed after waiting an additional 4-7 days.

#### Post-hoc visualization of viral labeling

as described in Michael et al. (2021).

#### Esr1 immunohistochemistry

Mice were deeply anaesthetized with isoflurane and then transcardially perfused with ice-cold 4% paraformaldehyde in 0.1 M phosphate buffer, pH 7.4 (4% PFA). Dissected brain samples were post-fixed overnight in 4% PFA at 4 °C, cryoprotected in a 30% sucrose solution in PBS at 4 °C for 48 hours, frozen in Tissue-Tek O.C.T. Compound (Sakura), and stored at –80 °C until sectioning. Brains were cut into 80 µm coronal sections, rinsed 3X in PBS, permeabilized for 3 hours in PBS containing 1% Triton X (PBST), and then blocked in 0.3% PBST containing 10% Blocking One (Nacalai Tesque; blocking solution). Sections were processed for 48 hours at 4 degrees with the primary antibody in blocking solution (1:1250, rabbit anti-Esr1, Invitrogen PA1-309), rinsed 3×10 minutes in PBS, then processed for 48 hours at 4 degrees with secondary antibodies in blocking solution (1:1000, Alexa Fluor 647 goat anti-rabbit, Jackson Laboratories, plus 1:500 NeuroTrace, Invitrogen). Tissue sections were then rinsed again 3×10 minutes in PBS, mounted on slides, and coverslipped using Fluoromount-G (Sothern Biotech). After drying, slides were imaged with a 10X or 20X objective on a Zeiss 700 laser scanning confocal microscope and the overlap between GFP+ and Esr1+ neurons was counted manually.

#### *In situ* hybridization using hybridization chain reaction (HCR)

as described in Michael et al. (2021).

#### Fiber photometry

On the day of testing, the implanted fiber optic cannula was plugged into a three-channel multi-fiber photometry system (Neurophotometrics, Ltd.). In this system, 470 nm (for imaging in green), 560 nm (for imaging in red), and 415 nm (control signal to detect calcium-independent artifacts) LEDs are bandpass filtered and directed down a fiber optic patch cord which is coupled to the implanted fiber optic cannula. Emitted GCaMP fluorescence is captured through the patch cord and split by a 532 dichroic, bandpass filtered, and focused onto opposite sides of a CMOS camera. Data were acquired using Bonsai software by drawing a region of interest (ROI) around the green image of the patch cord and calculating the mean pixel value. The blue and green channels were median filtered and the blue channel was fit to and subtracted from the green signal. The resulting calcium signal was analyzed using custom matlab codes.

#### Whole-cell recordings

as described in Michael et al. (2021).

#### Data availability

Data will be deposited to the Duke Research Data Repository. We will deposit 3 types of data in the repository: (1) confocal microscope images, (2) audio and video files from the mice used in this study and (3) slice electrophysiology data. All other data analyzed in this study are included in the manuscript and supporting files.

#### Code availability

Custom-written Matlab codes used in this study have been deposited to the Duke Research Data Repository, under the https://doi.org/10.7924/r4cz38d99 and are also available from the co-corresponding authors. The latest version of Autoencoded Vocal Analysis, the Python package used to generate, plot, and analyze latent features of mouse USVs, is freely available online: https://github.com/jackgoffinet/autoencoded-vocal-analysis.

### QUANTIFICATION AND STATISTICAL ANALYSES

#### Statistics

Parametric, two-sided statistical comparisons were used in all analyses unless otherwise noted (alpha=0.05). No statistical methods were used to predetermine sample sizes. Error bars represent standard error of the mean unless otherwise noted. Mice were selected at random for inclusion into either experimental or control groups for optogenetic experiments. Mice were only excluded from analysis in cases in which viral injections were not targeted accurately, or in cases with absent or poor viral expression.

## Acknowledgements

Thanks to Michael Booze for mouse husbandry, thanks to Katherine Tschida for help with HCR in situ hybridization, and thanks to Fan Wang’s lab for providing rabies tracing-related viruses. This work is supported by NIH grants DC 013826 (to RM) and MH 117778 (to FW and RM).

## Ethics

Animal experimentation: All experiments were conducted according to protocols approved by the Duke University Institutional Animal Care and Use Committee protocol (# A227-17-09).

## Author contributions

VM, SX and RM designed the experiments. SX and VM conducted the experiments. VM and SX analyzed data. SX, VM and RM wrote the manuscript, and all authors approved the final manuscript.

---

**Figure 1 —figure supplement 1:**
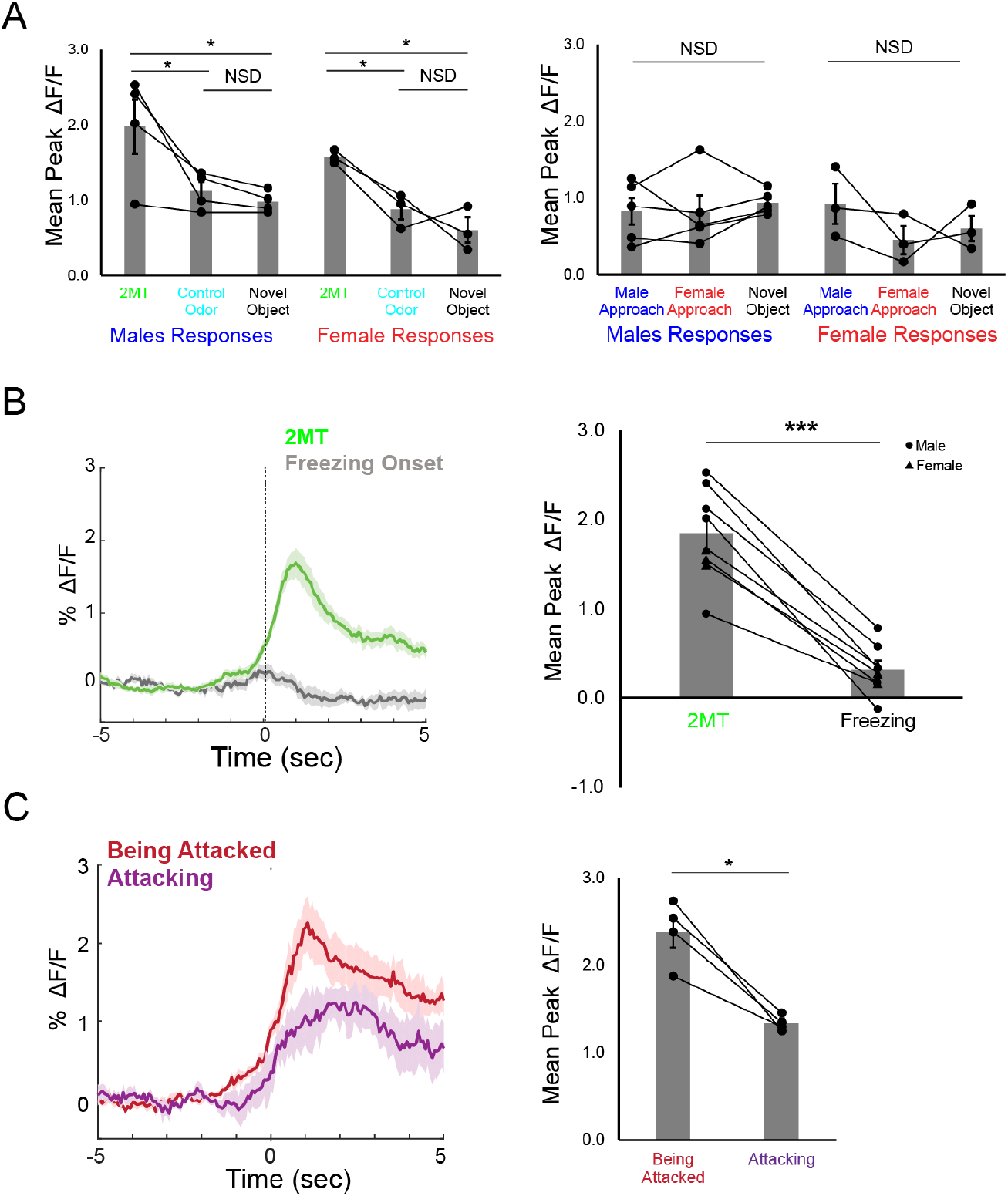
(A) Mean peak Amg_C/M-PAG_ activity in male and female mice as the animal approached and investigated a dish containing either 2MT, a control odor (ethyl tiglate) or a novel plastic “toy” (left; N=7 mice, 4 males, 3 females) or as the animal approached a conspecific male or a female (right; N=8 mice, 5 males, 3 females). (B) Average Amg_C/M-PAG_ activity (left) and mean peak Amg_C/M-PAG_ activity (right) as the animal approached and investigated a dish containing 2MT or at the onset of freeze in the presence of 2MT (N=8 mice). (C) Average Amg_C/M-PAG_ activity (left) and mean peak Amg_C/M-PAG_ activity (right) as the male attacked or got attacked (N=4 males).

## References

Akre, K. L., Farris, H. E., Lea, A. M., Page, R. A., & Ryan, M. J. (2011). Signal perception in frogs and bats and the evolution of mating signals. Science, 333(6043). https://doi.org/10.1126/science.1205623

Bhattacharjee, S., Kashyap, R., Abualait, T., Annabel Chen, S. H., Yoo, W. K., & Bashir, S. (2021). The Role of Primary Motor Cortex: More Than Movement Execution. In Journal of Motor Behavior (Vol. 53, Issue 2). https://doi.org/10.1080/00222895.2020.1738992

Breitfeld, T., Bruning, J. E. A., Inagaki, H., Takeuchi, Y., Kiyokawa, Y., & Fendt, M. (2015). Temporary inactivation of the anterior part of the bed nucleus of the stria terminalis blocks alarm pheromone-induced defensive behavior in rats. Frontiers in Neuroscience, 9(SEP). https://doi.org/10.3389/fnins.2015.00321

Callaway, E. M., & Luo, L. (2015). Monosynaptic circuit tracing with glycoprotein-deleted rabies viruses. Journal of Neuroscience, 35(24). https://doi.org/10.1523/JNEUROSCI.0409-15.2015

Chabout, J., Sarkar, A., Dunson, D. B., & Jarvis, E. D. (2015). Male mice song syntax depends on socia lcontexts and influences female preferences. Frontiers in Behavioral Neuroscience, 9(APR). https://doi.org/10.3389/fnbeh.2015.00076

Chen, J., Markowitz, J. E., Lilascharoen, V., Taylor, S., Sheurpukdi, P., Keller, J. A., Jensen, J. R., Lim, B. K., Datta, S. R., & Stowers, L. (2021). Flexible scaling and persistence of social vocal communication. Nature, 593(7857). https://doi.org/10.1038/s41586-021-03403-8

Day, H. E. W., Masini, C. V., & Campeau, S. (2004). The pattern of brain c-fos mRNA induced by a component of fox odor, 2,5-dihydro-2,4,5-Trimethylthiazoline (TMT), in rats, suggests both systemic and processive stress characteristics. Brain Research, 1025(1–2). https://doi.org/10.1016/j.brainres.2004.07.079

Gogolla, N. (2017). The insular cortex. In Current Biology (Vol. 27, Issue 12). https://doi.org/10.1016/j.cub.2017.05.010

Hashikawa, K., Hashikawa, Y., Tremblay, R., Zhang, J., Feng, J. E., Sabol, A., Piper, W. T., Lee, H., Rudy, B., & Lin, D. (2017). Esr1+ cells in the ventromedial hypothalamus control female aggression. Nature Neuroscience, 20(11). https://doi.org/10.1038/nn.4644

Jürgens, U. (2009). The Neural Control of Vocalization in Mammals: A Review. In Journal of Voice (Vol. 23, Issue 1). https://doi.org/10.1016/j.jvoice.2007.07.005

Jürgens, Uwe. (2002). Neural pathways underlying vocal control. In Neuroscience and Biobehavioral Reviews (Vol. 26, Issue 2). https://doi.org/10.1016/S0149-7634(01)00068-9

Karigo, T., Kennedy, A., Yang, B., Liu, M., Tai, D., Wahle, I. A., & Anderson, D. J. (2021). Distinct hypothalamic control of same- and opposite-sex mounting behaviour in mice. Nature, 589(7841). https://doi.org/10.1038/s41586-020-2995-0

Kimchi, T., Xu, J., & Dulac, C. (2007). A functional circuit underlying male sexual behaviour in the female mouse brain. Nature, 448(7157). https://doi.org/10.1038/nature06089

Kudwa, A. E., Michopoulos, V., Gatewood, J. D., & Rissman, E. F. (2006). Roles of estrogen receptors α and β in differentiation of mouse sexual behavior. Neuroscience, 138(3). https://doi.org/10.1016/j.neuroscience.2005.10.018

Lee, H., Kim, D. W., Remedios, R., Anthony, T. E., Chang, A., Madisen, L., Zeng, H., & Anderson, D. J. (2014). Scalable control of mounting and attack by Esr1+ neurons in the ventromedial hypothalamus. Nature, 509(7502). https://doi.org/10.1038/nature13169

Lin, D. Y., Shea, S. D., & Katz, L. C. (2006). Representation of Natural Stimuli in the Rodent Main Olfactory Bulb. Neuron, 50(6). https://doi.org/10.1016/j.neuron.2006.03.021

Maggio, J. C., & Whitney, G. (1985). Ultrasonic vocalizing by adult female mice (Mus musculus). Journal of Comparative Psychology (Washington, D.C. : 1983), 99(4). https://doi.org/10.1037/0735-7036.99.4.420

Michael, V., Goffinet, J., Pearson, J., Wang, F., Tschida, K., & Mooney, R. (2020). Circuit and synaptic organization of forebrain-to-midbrain pathways that promote and suppress vocalization. ELife, 9. https://doi.org/10.7554/ELIFE.63493

Moran, C. R., Gallagher, J. M., & Bridges, R. S. (2020). The role of the estrogen receptor-α gene, Esr1, in maternal-like behavior in juvenile female and male rats. Physiology and Behavior, 216. https://doi.org/10.1016/j.physbeh.2020.112797

Neunuebel, J. P., Taylor, A. L., Arthur, B. J., & Roian Egnor, S. E. (2015). Female mice ultrasonically interact with males during courtship displays. ELife, 4(MAY). https://doi.org/10.7554/eLife.06203

Paredes, R. G. (2003). Medial preoptic area/anterior hypothalamus and sexual motivation. In Scandinavian Journal of Psychology (Vol. 44, Issue 3). https://doi.org/10.1111/1467-9450.00337

Portfors, C. V. (2007). Types and functions of ultrasonic vocalizations in laboratory rats and mice. In Journal of the American Association for Laboratory Animal Science (Vol. 46, Issue 1).

Roberts, J. A., Taylor, P. W., & Uetz, G. W. (2007). Consequences of complex signaling: Predator detection of multimodal cues. Behavioral Ecology, 18(1). https://doi.org/10.1093/beheco/arl079

Root, C. M., Denny, C. A., Hen, R., & Axel, R. (2014). The participation of cortical amygdala in innate, odour-driven behaviour. Nature, 515(7526). https://doi.org/10.1038/nature13897

Salgado, S., & Kaplitt, M. G. (2015). The nucleus accumbens: A comprehensive review. In Stereotactic and Functional Neurosurgery (Vol. 93, Issue 2). https://doi.org/10.1159/000368279

Seyfarth, R. M., & Cheney, D. L. (2010). Production, usage, and comprehension in animal vocalizations. Brain and Language, 115(1). https://doi.org/10.1016/j.bandl.2009.10.003

Shipley, M. T., & Ennis, M. (1996). Functional organization of olfactory system. In Journal of Neurobiology (Vol. 30, Issue 1). https://doi.org/10.1002/(SICI)1097-4695(199605)30:1<123::AID-NEU11>3.0.CO;2-N

Tschida, K., Michael, V., Takatoh, J., Han, B. X., Zhao, S., Sakurai, K., Mooney, R., & Wang, F. (2019). A Specialized Neural Circuit Gates Social Vocalizations in the Mouse. Neuron, 103(3). https://doi.org/10.1016/j.neuron.2019.05.025

Warren, M. R., Clein, R. S., Spurrier, M. S., Roth, E. D., & Neunuebel, J. P. (2020). Ultrashort-range, high-frequency communication by female mice shapes social interactions. Scientific Reports, 10(1). https://doi.org/10.1038/s41598-020-59418-0

Warren, Megan R., Spurrier, M. S., Roth, E. D., & Neunuebel, J. P. (2018). Sex differences in vocal communication of freely interacting adult mice depend upon behavioral context. PLoS ONE, 13(9). https://doi.org/10.1371/journal.pone.0204527

Zatorre, R. J., Belin, P., & Penhune, V. B. (2002). Structure and function of auditory cortex: Music and speech. In Trends in Cognitive Sciences (Vol. 6, Issue 1). https://doi.org/10.1016/S1364-6613(00)01816-7

Zhao, X., Ziobro, P., Pranic, N. M., Chu, S., Rabinovich, S., Chan, W., Zhao, J., Kornbrek, C., He, Z., & Tschida, K. A. (2021). Sex- And context-dependent effects of acute isolation on vocal and non-vocal social behaviors in mice. PLoS ONE, 16(9 September). https://doi.org/10.1371/journal.pone.0255640

